# A microphysiological system of sterile injury demonstrates neutrophil reverse migration via macrophage-derived extracellular vesicle crosstalk

**DOI:** 10.1101/2024.12.30.630550

**Authors:** Kehinde Adebayo Babatunde, Babatunde Fatimat Oluwadamilola, Adeel Ahmed, Wilmara Salgado-Pabon, David J Beebe, Sheena C Kerr

## Abstract

Persistent neutrophilic inflammation can lead to tissue damage and chronic inflammation, contributing to non-healing wounds. The resolution phase of neutrophilic inflammation is critical to preventing tissue damage, as observed in diseases characterized by influx of neutrophils such as atherosclerosis and non-healing wounds. Animal models have provided insight into resolution of neutrophilic inflammation via efferocytosis and reverse migration (rM); however, species-specific differences and complexity of innate immune responses make translation to humans challenging. Thus, there is a need for *in vitro* systems that can elucidate mechanisms of resolution of human neutrophilic inflammation. Here, we developed a human microphysiological system (MPS) to mimic an inflammatory sterile injury (SI) microenvironment to study the role of macrophage derived extracellular vesicles (M-EVs) in determining the resolution of inflammation via neutrophil rM. The MPS integrates a human umbilical vein endothelial cell (HUVEC) lined lumen, injury site spheroid, human neutrophils, macrophages and macrophage derived EVs to investigate the role of M-EVs in neutrophil rM *in vitro*. The key features of the MPS enabled us to demonstrate that EVs derived from macrophage subsets modulate migratory behavior in primary neutrophils differently in specific inflammatory microenvironments. Importantly, we identified a new mechanism underlying neutrophil rM via M-EV, where neutrophils exposed to M2-EV-derived IL-8 migrate away from the SI site upon reaching the site, using the SI MPS. Overall, our SI MPS system demonstrates a reverse migratory pattern in human primary neutrophils, advancing the study of the resolution of inflammation via M-EVs.

## INTRODUCTION

Sterile injury (SI) and non-sterile injury (nSI) are broad terms covering many inflammatory insults that occur in the absence and presence of microorganisms respectively (*1*). Neutrophils and macrophages are recruited within the first few hours of injury, with neutrophils usually considered the first responders, that sense chemotactic signals and migrate rapidly to sites of infection and injury during the innate immune response (*2*). However, persistent neutrophilic inflammation can lead to tissue damage and chronic inflammation contributing to non-healing wounds. The resolution phase of neutrophil inflammation is critical to preventing tissue damage, as observed in diseases characterized by an influx of neutrophils such as non-healing wounds, atherosclerosis and arthritis (*3*). Two main mechanisms involving neutrophil-macrophage crosstalk have been proposed to promote the resolution of inflammation during tissue repair, with both involving neutrophil–macrophage crosstalk: efferocytosis (apoptotic cells are removed by phagocytic cells) and reverse migration (rM) (neutrophils moving away from the SI site). While neutrophil-macrophage crosstalk has been implicated in both mechanisms in animal models (*3–6*), data is lacking for human cells.

Macrophages play a key role in neutrophil recruitment to tissues by recognizing damage-associated molecules. Pro-inflammatory mediators released from macrophages recognize several receptors on neutrophils and direct neutrophil recruitment via chemokines (*7*). Importantly, macrophage-neutrophil crosstalk has been implicated in the resolution of inflammation (*8*). Several modes of cell-to-cell communication have been identified including the release of soluble signaling factors, direct cell-to-cell contact, and signaling cargoes transferred via extracellular vesicles (EVs). EVs are small vesicles released by almost all eukaryotic cell types and are surrounded by lipid membranes. They are involved in various cellular functions that transmit information between cells, for instance, EVs transfer protective or proinflammatory information (*9*). Chemokines are enclosed in EVs, and they can modulate cellular chemotaxis. Monocytes/macrophages may be recruited for migration in atherosclerosis by monocyte-derived exosomes containing inflammatory factors(*10*) thus showing that EVs play an important role in migration. However, whether EVs, such as those secreted by macrophages, determine resolution of inflammation via primary neutrophils rM remains to be studied.

Previous studies have employed straight microfluidic channels to evaluate primary leukocyte migration, including neutrophils, migrating away from a chemoattractant source (*11*, *12*). Using advanced 3D bioprinting techniques, straight microfluidic channels with established chemoattractant gradients were used to study primary leukocyte migration (*11*). These systems were able to demonstrate different leukocyte migration signatures including rM with exogenous chemoattractant sources. However, these systems do not incorporate physiological components such as endothelial cell lined lumens to mimic vasculature, endogenous inflammatory source (self-generating) and biological components such as EVs that are shed by immune cells during inflammation.

Microphysiological systems (MPS) enable the integration of three-dimensional (3D) complexity, spatial organization, and relevant cell-cell and cell-extracellular matrix (ECM) interactions that are critical for modeling *in vivo*–like responses such as inflammation (*13*). Immune cell trafficking across the endothelial barrier has been studied using MPS (*14*). Interestingly, MPS can include the key elements likely to be important for studying rM such as integration of immune cell components, source of focal inflammatory signals, and injury site architecture that can modulate the resolution of inflammation via primary neutrophil rM *in vitro*.

In this work, we developed an MPS of nSI and SI models. Using the SI model, we investigated how M-EVs could modulate rM in primary neutrophils *in vitro*. By using a micro-molding technique to create a luminal structure out of a collagen hydrogel, we lined the lumens with HUVECs to recapitulate the cell components and geometries of blood vessels with an incorporated spheroidal injury site, mimicking a focal inflammatory source. This micro-engineered MPS of a human blood vessel and inflammatory SI site, provides a means for studying rM in primary neutrophils as a result of their interaction with M-EVs. The MPS ensures that neutrophils in the HUVECs lined lumen are located in close proximity to the SI site, thus enabling identification and tracking of specific neutrophil migration signatures (rM) in response to the SI.

Using the MPS, we demonstrated that neutrophils exposed to pro-inflammatory macrophage derived EVs (M1-EV) showed persistent migration toward the SI site however, anti-inflammatory macrophage EVs (M2-EV) via interleukin-8 (IL-8) induce neutrophils to migrate away from the SI site once they reach the site. The data presented here demonstrate a new M-EV related mechanism underlying rM in primary neutrophils not previously well defined *in vitro* using traditional techniques or other microfluidic platforms. In addition, our data suggest that M-EVs could potentially be important for the design of anti-inflammatory agents.

## MATERIALS AND METHODS

### Focal injury MPS development

The focal injury LumeNEXT MPS was fabricated using micro-molding techniques to generate luminal structures with an ECM scaffold as previously described by Jimenez-Torres et al (*15*). The MPS comprises a single lumen, which was equipped with an inlet and outlet port, within a single chamber with perpendicularly oriented side ports. Two stacked polydimethylsiloxane (PDMS) layers, with microscale features patterned into them, formed the culture chambers in which scaffolding ECM gel can be loaded, while a removable PDMS rod formed the hollow lumen structure surrounded by the ECM gel. Briefly. the molds for the PDMS layers were made using SU-8 100 (Microchem), which were spin-coated onto wafers, soft-baked (i.e., heat at 65°C for 30 to 40 min and then at 95°C for 90 min depending on layer thickness), exposed to ultraviolet (UV) through a mask of desired patterns, and post-baked at 95°C for 20 to 30 min. This procedure was repeated for additional layers before development in propylene glycol monomethyl ether acetate (MilliporeSigma). After developing, PDMS (Sylgard 184 Silicon Elastomer Kit, Dow Corning Corporation) was applied to the masters at a ratio of 10:1 base to curing agent and allowed to polymerize for 4 hours at 80°C. The rods were drawn from needles with gauge size, 23G (340-μm inner diameter). Before device assembly, the PDMS layers and rods were soaked in 100% ethanol for several days to extract any uncured PDMS oligomers. Following PDMS extraction, the rods were placed in between two layers, across the body of the chamber (3 mm in length) in ledge features stemming from the smaller inlet and larger outlet ports to hold the rods in the middle of the chamber. The side ports (4 mm apart) of the chamber were used to fill the chamber with ECM gel, and the height of the chamber was about 1.25 mm. Once assembled, the PDMS layers were oxygen-plasma–bonded onto a glass-bottom MatTek dish using a Diener Electronic Femto Plasma Surface System.

### ECM gel preparation/loading and spheroid formation

The bonded devices were UV-sterilized for 20 min and moved to the biosafety cabinet before ECM gel loading. To promote matrix adhesion to PDMS, the device chambers were treated with 1% polyethylenimine (MilliporeSigma) in deionized (DI) water solution for 10 min, followed by a 30-min treatment of 0.1% glutaraldehyde (MilliporeSigma) in DI water solution. Following surface treatment, devices were flushed with DI water solution five times to remove excess glutaraldehyde. A high concentration rat tail collagen I (Col-I) (Corning-8.08 mg/ml-354249, *Corning*, NY, USA) neutralized with 0.5 N sodium hydroxide (Thermo Fisher Scientific) was mixed with 7.5 pH 5× phosphate-buffered saline (PBS), complete growth medium to achieve a final ECM solution containing Col-I (3 mg/ml). The pH of the ECM mix was adjusted to pH 7.2 before loading into the gel chamber of the device. The devices were first kept at room temperature for 20 min and then moved to an incubator at 37°C for at least 1 hour before cell loading. To prevent dehydration during polymerization, PBS was added to the MatTek dish surrounding the devices. PDMS rods were then removed, leaving behind luminal structures within the ECM gel that can be lined with cells.

Human derived dermal fibroblast spheroids were generated using the hanging drop method with methylcellulose (*16*). Briefly, 12 g of high viscosity methylcellulose (Sigma, M0512) were dissolved in one liter of RPMI1640, and centrifuged at 4000 g for 2.5 h. Only the clear supernatant was used in the next step. Cell suspension (10^5^ cells/ml) and this methylcellulose solution were mixed in a 4:1 v/v ratio, and 25 μl droplets (2×10^3^ cells) were placed on the lid of Petri dishes. Sterile water was placed on the bottom of the dishes to prevent evaporation from droplets, the lids were replaced, and the Petri dishes incubated at 37°C and 5% CO_2_. After 1 day, a single well-defined spheroid was generated per drop. Five to ten spheroids are mixed with the prepared ECM gel and one or two spheroid(s) are gently picked and loaded with the ECM into the gel chamber of the device.

### HUVEC lumen formation

HUVECs (ATCC) were maintained in EGM-2 MV BulletKit medium (Lonza) and used until passage 8. To generate endothelial vessels, HUVECs were detached using a trypsin/EDTA solution and resuspended at 15 million cells/ml of EGM-2 MV. Three microliters of endothelial cell suspension were introduced into the cell-free lumen next to the spheroid injury site, and the device was rotated every 30 min over a 1-hour period as previously described by Jimenez-Torres et al (*15*). EGM-2 MV media was added to the gel ports and epithelial inlet/outlet ports during this process to nourish the HUVEC cells. After lining the tube with endothelial cells, nonadherent cells were gently washed with EGM-2 MV media.

### Isolation of human neutrophils

Neutrophils were isolated from peripheral human blood acquired by venipuncture from healthy donors under an approved IRB protocol at UW Madison. Isolation was performed within 1 h after the blood draw using the Easy Sep human neutrophil negative isolation kit (STEMcell Technologies, Vancouver, Canada) following the manufacturer’s protocol. After isolation, neutrophils were suspended in RPMI 1640 media containing 10% FBS (Thermo Fisher Scientific) and 1% penicillin/streptomycin.

### Isolation of human monocytes

Monocytes were isolated from peripheral human blood acquired by venipuncture from healthy donors under an approved IRB protocol at UW Madison. Isolation was performed within 1 h after the blood draw using the Easy Sep direct human monocyte isolation kit (STEMcell Technologies, Vancouver, Canada) following the manufacturer’s protocol. After isolation, monocytes were cultured and differentiated into macrophages in DMEM media containing 10% FBS (Thermo Fisher Scientific) and 1% penicillin/streptomycin as described below.

### Human monocytic cell line (THP-1) culture, differentiation and polarization

Human monocytic THP-1 cells were maintained in culture in Roswell Park Memorial Institute medium (RPMI 1640, Invitrogen) culture medium containing 10 % of heat inactivated fetal bovine serum (Invitrogen). THP-1 monocytes are differentiated into macrophages by 24 h incubation with 150 nM phorbol 12-myristate 13-acetate (PMA, Sigma, P8139) followed by 24-48 h incubation in fresh RPMI medium. Macrophages were then polarized to M1 macrophages by incubation with 20 ng/ml of IFN-γ (R&D system) and 20 pg/ml of LPS (Sigma) for 48 hours. Macrophage M2 polarization was obtained by incubation with 20 ng/ml of interleukin 4 (peproTech, 200-04-20UG) and 20 ng/ml of interleukin 13 (peproTech, 200-13-10UG).

To confirm M1 and M2 phenotypes, we carried out immunofluorescence assay by fixing cell samples using 4% Paraformaldehyde for 5 minutes and permeabilizing in 0.1% triton-X for 10 minutes. Blocking was done in 3% Bovine serum albumin for 1h at room temperature. For M1 phenotype staining, we stained with 5µg/ml of CD 80 primary antibody (MA5-15512-Thermo fisher) overnight at 4 degree Celsius and detection was achieved by staining with an anti-mouse Alexa fluor 547 secondary antibody (Thermo fisher) for 1h at room temperature and Hoechst to detect nuclei. For M2 phenotype staining we stained with FITC conjugated CD 163 antibody (FabGennix) overnight at 4 degree Celsius and Hoechst to detect nuclei. Images were taken using a Nikon spinning disk confocal microscope.

### Primary monocyte differentiation & polarization and M-EV isolation

Primary monocytes were incubated in DMEM and differentiated into macrophages for 6 days in the presence of 100 ng/ml M-CSF and GM-CSF. Next, we polarized them into either M1 or M2 by treating with LPS (20pg/ml) or IL-4 and IL-13 (20ng/ml) respectively for 48 hours. The conditioned medium was collected and centrifuged at 1,000 × *g* for 20 min to remove cell debris, followed by centrifugation at 2,000 × *g* for 20 min and 10,000 × *g* for 30 min at 4°C. The supernatant was subsequently collected and filtered passed through 0.22-μm filters (MilliporeSigma) and centrifuged in Beckman Coulter Optima L-80XP at 100,000 × *g* for 70 min at 4°C to pellet EVs. Next, the EV pellets were washed with sterile PBS and subjected to another cycle of ultracentrifugation at 100,000 × *g* for 70 min at 4°C. Finally, the pelleted EVs were carefully reconstituted in sterile PBS or lysed in Triton-X for western blot or RIPA buffer (Thermo Fisher Scientific) for the chemokine protein array. For all experiments, freshly isolated M-EVs were used or isolated a day before the experiment. For each experiment, neutrophils were incubated with 100 μg of isolated M-EVs. M-EVs concentration were measured using a nanodrop (Thermo Fisher).

### M-EV internalization and ICAM-1 immunofluorescent staining

Isolated M-EVs were labelled with PKH67 green fluorescent labelling kit. Briefly, M-EVs were fluorescently labeled with 20 μM PKH-67 green fluorescent labeling kit (MINIKIT-67, Sigma-Aldrich) for 5-6 minutes at room temperature (protected from light), washed three times and incubated with neutrophils for 4h (50,000 neutrophils: 100µg M-EVs) before washing cells three times to remove unbound EVs. For uptake inhibition experiments neutrophils were co-incubated with different compounds for 30-45 mins and excess drugs were washed off three times using PBS. Cells were fixed in 4% Paraformaldehyde for 5 minutes and then permeabilized in 0.1% triton-X for 5 minutes. Blocking was done in 3% Bovine serum albumin for 1h at room temperature. Next, we stained for F-actin with phalloidin-Texas red (Invitrogen) and with the nuclear dye Hoechst (Thermo-Fisher). Images were taken using a confocal Nikon Microscope.

To confirm reverse migrated neutrophil phenotype in our MPS during SI, the whole MPS device was fixed using 4% Paraformaldehyde for 5 minutes and permeabilized in 0.1% triton-X for 10 minutes. Blocking was done in 3% Bovine serum albumin for 1h at room temperature. For ICAM-1 stanning-the device was stained with primary antibody (Thermo fisher) overnight at 4 degree Celsius and detection was achieved by staining with an Alexa fluor 488 conjugated secondary antibody (Thermo fisher) for 1h at room temperature and Hoechst to detect nuclei. Images were taken using a Nikon Ti Eclipse microscope. Quantification of ICAM-1 staining around the sterile injury area was done by analyzing immunofluorescent images using the analyze particle plugin on ImageJ.

### Chemical inhibitors

For inhibiting the uptake pathways in neutrophils, we included Colchicine (Sigma-Aldrich) in the media at a concentration of 31 nM, Latrunculin (Cayman Chemicals) at a concentration of 5 µM, nocodazole (Cayman Chemicals) at a concentration of 20 µM and dynasore at a concentration of 80 µM. All chemical inhibitors were accompanied and compared to the appropriate vehicle control.

### Flow cytometry

Following neutrophils and M-EVs incubation, cells were harvested and washed three times in PBS to remove excess unbound M-EVs. Prior to incubation with neutrophils, M-EVs were stained with PKH-67 fluorescent dye for visualization as described above. After harvest and washing, cells were re suspended in FACS buffer (0.1% BSA in PBS and 2% FBS). Images were taken using the Image Stream cytometer to confirm M-EVs uptake.

### Bacteria culture

*Staphylococcus aureus* tagged with green fluorescent protein was a gift from JD Sauer, Department of Microbiology, University of Wisconsin Madison. Briefly, bacteria were maintained as a −80°C stock and streaked onto Luria-Bertani agar (LB; 2% agar) plates 2 days before each experiment. Single colonies of *S. aureus* in LB broth were grown overnight from these plates at 37°C with 180 rpm of shaking a day before experiment. For the bacteria ring spheroid generation, a MOI of 10:1 (MOI, bacterium: eukaryotic cell ratio). was used to infect the spheroid and cultured overnight to allow bacteria to infect the spheroid.

### Colony Forming Unit Assay

*Staphylococcus aureus* GFP tagged were cultured overnight as described above. The next day, the bacteria were diluted 1:20 and let grown to the exponential phase (OD=600) within 2 hours. The bacteria were opsonized for 10 min in human serum. Plates were incubated at 37°C and 5%CO_2_ for 20 min with neutrophils prior to the addition of *S. aureus* at a multiplicity of infection 10:1 (MOI, bacterium: eukaryotic cell ratio). After the incubation time (3-4 hours), the medium was removed and replaced with medium containing gentamicin (10 μg/mL in RPMI) for 30 min to kill extracellular bacteria. The cells were washed and lysed in ice-cold distilled water containing 1% Triton X-100 for 1 hour for lysis of neutrophils and were plated on LB agar, incubated for overnight at 37°C and viable bacteria colonies were quantified using Image J. Growth experiments were repeated at least with three independent donors.

### Nanoparticle tracking analysis

Particle size and concentration distribution of the isolated M-EVs were measured using nanoparticle tracking analysis. The nano tracking was performed at the pharmacy department, University of Wisconsin Madison. Briefly, the M-EV samples were diluted 1:5000 in filtered sterile PBS. Each sample analysis was conducted for 60s and measured three times using the automatic analysis settings of the NanoSight instrument.

### Transmission electron microscopy (TEM)

The TEM was performed at the School of Medicine and Public Health, University of Wisconsin Madison. Briefly, pelleted EVs from either primary macrophages or THP-1 cells were negatively stained with Nano-W (Nanoprobes; Yaphank, NY) using the two-step method. A 2 µl droplet of samples was placed on a formvar coated 300 mesh Cu Thin-Bar grid (EMS; Hatfield, PA), coating side down. The excess was wicked with a wedge of filter paper and allowed to dry. Immediately after the droplet dried, 2 μl droplet of Nano-W was applied, wicked again with clean new filter paper, and allowed to dry. The sample was viewed on a Philips CM120 transmission electron microscope at 80 kV and documented with an AMT BioSprint12 (Woburn MA) digital camera.

### Chemokine protein array

Custom human cytokine arrays (AAH-CHE-1-4; RayBio; RayBiotech, Norcross, GA) were used following the manufacturers protocols. Primary monocyte derived macrophage EV lysates or THP-1 derived macrophage lysates were incubated with membranes coated with anti-human chemokine Abs. Briefly, the membranes were incubated with blocking buffer at room temperature for 30 min and incubated with each sample at room temperature for 90 min. The membranes were washed three times with Wash Buffer I and twice with Wash Buffer II at room temperature for 5 min per wash. The membranes were then incubated with biotin-conjugated antibody at room temperature for 90 min. Finally, the membranes were washed and incubated with HRP-conjugated streptavidin at room temperature for 2 h and with detection buffer for 2 min. We used an iBright Imager (ThermoFisher, Invitrogen) for detection.

### Generation of IL-8 KO THP-1 cell line using CRISPR

A CRISPR kit targeting the first exon of IL-8 gene was obtained from Origene (cat. no. KN402075). Parental THP-1 cells were co-transfected with a gRNA vector plasmid targeting sequences 5’-TGTAAACATGACTTCCAAGC-3’ or 5’-CAAGAGAGCCACGGCCAGCT-3’ and a donor vector (LoxP-EF1A-tGFP-P2A-Puro-LoxP (2739 bp)) containing a selection cassette expressing GFP–puromycin was inserted into the targeted site in human IL-8 gene. Cells were transfected using the a Nucleofector (Lonza) following the manufacturer’s suggested conditions. Selected cells (GFP^+^ and puromycin resistant) were serially diluted to form single cell-derived colonies. The colonies were further screened for expression of IL-8 by immunoblotting, chemokine protein array.

### Migration assay

Migration assay was carried out using the LumeNEXT MPS. Briefly, about 50000 THP-1 WT or IL-8 KO THP-1 cells were embedded in the ECM of the LumeNEXT MPS. Next, primary neutrophils were loaded into the HUVEC seeded lumen and the device was incubated at 37°C and 5%CO_2_ overnight to study migratory response. Prior to loading neutrophils into the lumen, the embedded THP-1 cells were incubated in the collagen for about 3-4 hours. Images of the devices were taken using the Nikon Ti Eclipse microscope after overnight incubation and images were analyzed using the threshold and analyze particle plugin in image J.

### Western Blot

The total protein was extracted from isolated EVs or THP-1 cells using RIPA lysis buffer or 1% Triton-X100 respectively. Equal concentrations of protein extracted from 100µg of EVs were separated on 12% SDS-PAGE gels and transferred onto PVDF membranes. The blots were subsequently blocked for 1 h with 3% BSA. The EV protein blots were incubated with primary Abs (1:1000) against CD63 and CD81 (Thermo Fisher Scientific). The THP-1 protein blots were incubated with primary antibody against IL-8 and Beta-Actin (1:1000) (Thermo Fisher Scientific). Next, the blots were washed three times to remove excess antibody. For IL-8 KO THP-1, membranes were incubated with fluorescence-conjugated anti-rabbit for 1h at room temperature and images were taken using an azure biosystems imaging system. For the EV membrane, membranes were incubated with HRP anti-mouse Abs (LI-COR Biosciences) for 1 h at room temperature. CD63 and CD81 EV proteins were visualized using an ECL kit (Thermo Fisher Scientific) and images were taken an azure biosystems imaging system.

### Analysis and quantification of reverse migrated neutrophils

Acquired images and videos were exported and deconvoluted using Leica super STED microscope software. Exported imaged were imported to ImageJ software package (NIH) for analysis. To assess *in vitro* neutrophil migration behaviors, at the least three independent 1024 × 1024 fields of view (FOV) around the edge of the injury were imaged for 3-4-hours post spheroid injury site set up. 5-15 neutrophils per FOV were randomly selected for tracking. Next, we generated the x and y coordinates, distance and velocity of each individual cell by using the manual tracking plugin on image J software. The following analysis were done using the generated coordinates:

### Image Alignment

The recorded plots were plotted using the Plotly package in Python. To simplify analysis, representation and comparison, all coordinates in the image were transformed such that the spheroid would be at (0,0). The following transformations were used:

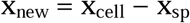

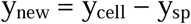

where (x_new_, y_new_) = new cell coordinates, x_sp_, y_sp_ = spheroid center as measured from the image.

### Plotting migration metrics

Distance was calculated between each frame using the distance formula:

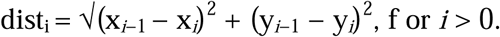

Next, velocity was calculated as dist_i_/time. To plot the cell location heatmap, kernel density estimation was employed with a gaussian kernel. Briefly, the density estimates for a given point on the plot is given by 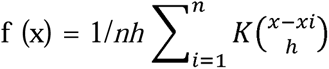 where, x*i* are the data points, K is a gaussian distribution, h is a bandwidth parameter, n is the number of datapoints. The SciPy package in python was employed to calculate the KDE. Bandwidth was calculated using Scott’s rule to faithfully maintain structural features of the data distribution and prevent oversampling.

### Identifying reverse migrated cells

Reverse migrated cells were defined as cells that came within 1 radius, radial distance of the spheroid center. These cells were identified by an x, y coordinate in their track where:

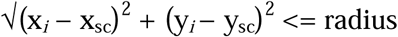

where x*_i_*, y*_i_* are cell coordinates at slice *i*, and x_sc_, y_sc_ are the coordinates for spheroid center. Since the spheroids are centered at (0, 0), the formula is reduced to:

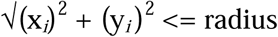

Following the selection, the track is divided into three segments, the approach segment includes all (x_a_, y_a_) coordinates before the cell entered the threshold region, (x_s_, y_s_) are the coordinates for when the cell is in the spheroid region, and (x_r_, y_r_) are all the points downstream after the cells exits the threshold region the for the last time to reverse migrate i.e. the cell did not re-enter the spheroid again.

### Calculating track straightness

To calculate track straightness, first the total distance d_total_ travelled by each cell was calculated for each migratory segment. Next, the displacement *disp* of the cell was calculated for each segment by using the distance formula on the first x*_s,i,_* y*_s,i_* and last coordinate x*_s,n,_* y*_s,n_* of a given x*s*. Finally, the straightness was defined as the ratio of displacement to distance *str* = *disp*/d_total_. A value of 1 indicated complete straightness.

### Statistical analysis

Statistical significance of the differences between multiple data groups were tested using two-way Analysis of Variance (ANOVA) in GraphPad Prism (GraphPad Software, version 10.3.1). Within ANOVA, significance between two sets of data was further analyzed using two-tailed t-tests. Non-paired Student’s t-test was performed for determining statistical significance between two conditions. All error bars indicate standard errors of means (SEM). ns, not significant (*p* ≥ 0.05); \**p*< 0.05; \*\**p* < 0.01; \*\*\**p* < 0.001.

## RESULTS

### Focal sterile injury MPS for demonstrating rM in primary neutrophils

rM has been described in neutrophils using animal models (*4*, *17*). However, evidence of rM in human neutrophils are still lacking, which is partly due to the unavailability of *in vitro* platforms that mimic the *in vivo* microenvironment for studying this process in human neutrophils. Importantly, such a platform must create an enabling microenvironment that includes primary immune cell components, a focal inflammatory signal source and injury site architecture that can demonstrate neutrophil rM in *vitro*.

We designed an MPS, based on the LumeNEXT platform (*15*), that incorporates an endothelial microvessel, an endogenous inflammatory signal [Damage associated molecular patterns (DAMPs), pathogen associated molecular patterns (PAMPs)] source, primary immune cell derived EVs and primary immune cells to allow the analysis of host immune responses during both SI (Fig. 1A) and nSI (Fig. 1B) in a physiologically relevant culture microenvironment. We generated lumen structures within a collagen-based ECM hydrogel using a micro-molding technique for modeling luminal geometries of blood vessel endothelium. The hydrogels are integrated into a PDMS elastomeric device with a central chamber, which contains the ECM gel, connected to two pairs of inlet and outlet ports that support PDMS rods used for molding the luminal structures (Fig. 1Ci). Following ECM gel polymerization, the rods are pulled out of the chamber, leaving behind hollow lumen structures (Fig. 1Cii).

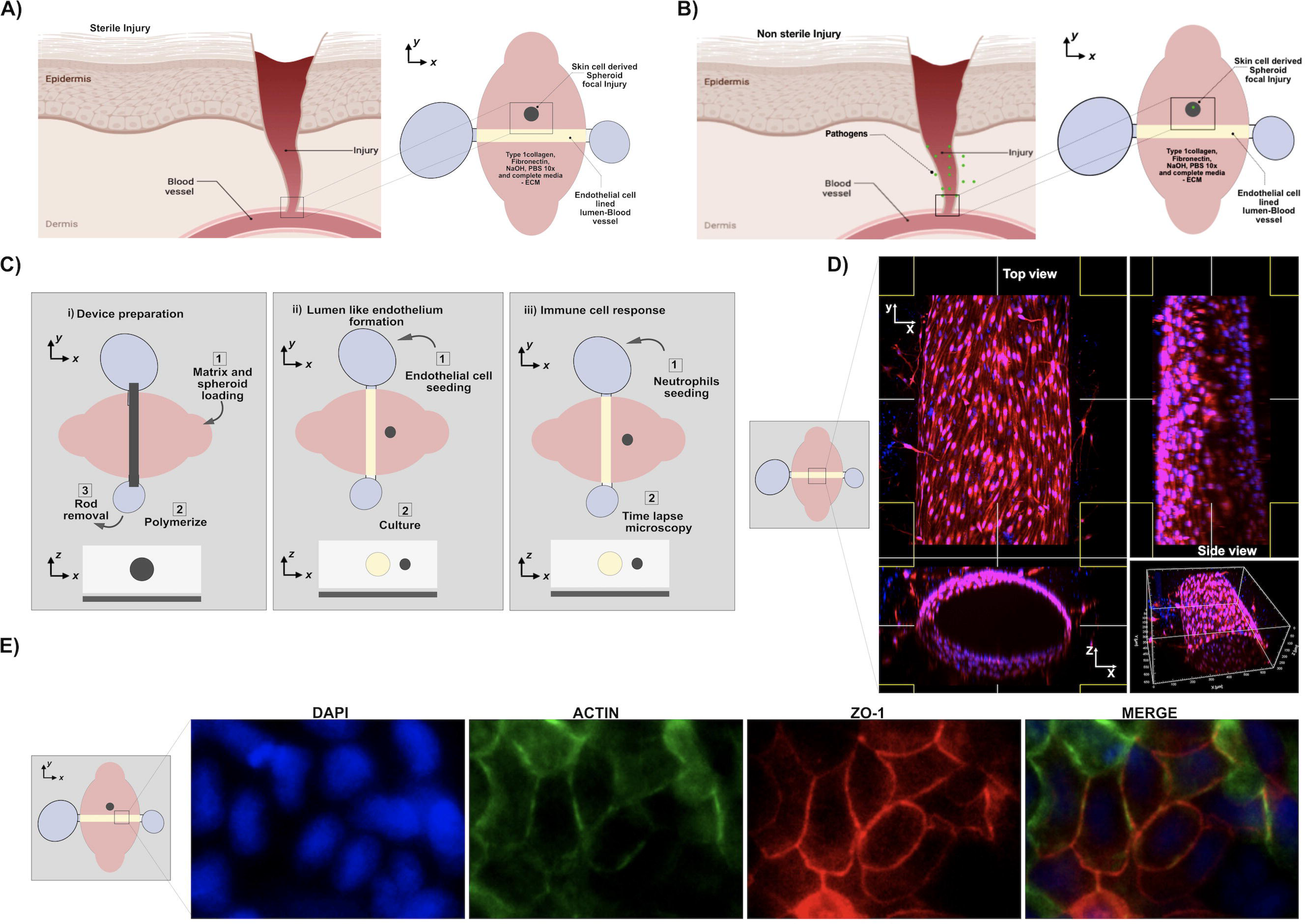
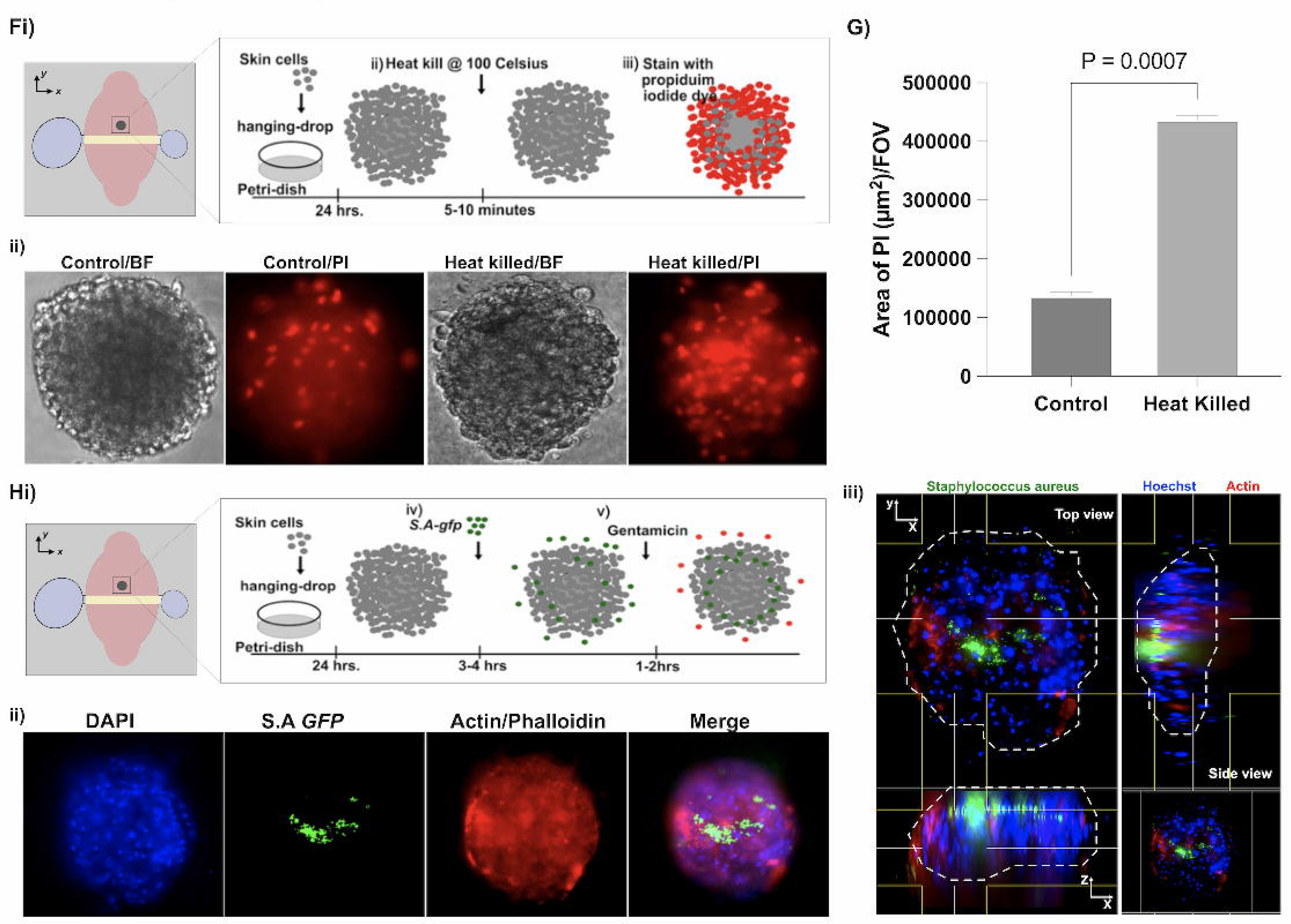

The inlet and outlet ports connected to the chamber provide direct access to the lumens, which can be used for cell seeding, addition of neutrophils and for the supply of medium during culture (Fig. 1Ciii). The endothelial vessel in the MPS was stained for calcein (live stain) (Fig. 1D) and for actin and for the tight junction marker ZO-1, which is involved in barrier function (Fig. 1E) (Supplementary Video 1), showing a fully cell-covered microvessel with tight junction formation. Next, we aimed to create a self-generating and endogenous inflammatory signal source adjacent to the endothelial cell lined lumen capable of establishing an inflammatory signal gradient between the lumen and the source within the MPS. To do this, we generated primary dermal fibroblast spheroids that will be modified to serve as the source of inflammatory signals (Fig. 1Fi) and mimic skin cell damage or infection.

To model SI focal sites, we starved (cultured in media without serum) the dermal cells for 24 hours prior to generating the spheroids and then heat killed them at 100°C for 5-10 mins (Fig. 1Fi-ii) with an increase in damaged/dead cells confirmed by propidium (PI) staining via microscopy (Fig. 1Fii & G). To model focal nSI in our MPS, we infected the dermal fibroblast spheroids with *Staphylococcus aureus* tagged with green fluorescent protein (*S.A^gfp^*) for 3-4 hours to form a bacteria ring around the spheroid (Fig. 1Hi-ii). Bacteria infiltrate the spheroid within 3-4 hours and extra-spheroidal bacteria were killed using a poorly diffusible antibiotic gentamicin (Fig. 1Hi-iii). The multiplicity of infection (MOI) in the nSI model was 1:10 of PMN to *S.Agfp*. *S.Agfp* at their log phase to generate the bacteria-spheroid. To incorporate the nSI (bacteria ring spheroid) or SI (heat killed spheroid) in the MPS, these spheroids were added adjacent to the endothelial lumen in a 3D extracellular matrix in our MPS device at the time of matrix addition and confirmed via microscopy (Fig. 1Ci) (Supplementary Video 2).

Together, our models mimic DAMPs and PAMPs signals generated by an endogenous self-generating spheroidal source present in an inflammatory microenvironment adjacent blood vessel, and immune cell components that could provide insight into neutrophil macrophage interaction during injury in a more physiologically relevant context.

### Isolation and characterization of macrophage derived extracellular vesicles

There are several modes of communication by which cells communicate with each other which include the release of soluble signaling factors, cell-to-cell contact, and signaling cargoes transferred via EVs. Macrophage-neutrophil crosstalk has been implicated in resolution of inflammation. Macrophages are polarized on a spectrum between pro-inflammatory (M1) or anti-inflammatory phenotypes (M2). To study the modulatory effect of M1-EVs and M2-EVs on neutrophils during SI and nSI injury *ex vivo*, we first differentiated primary monocytes into macrophages by treating with GM-CSF and M-CSF for 6 days and then polarized primary macrophages into either M1 or M2 by treating with LPS or IL4/IL13 respectively in a serum free media (*18*)(Supplementary fig. 1A). We then collected the culture supernatants from LPS-stimulated (M1) and IL4/1L-13 (M2) primary macrophages and isolated M-EVs by ultracentrifugation (*19*, *20*).

Next, we stained the lipid bi-layer of the isolated M-EVs with PKH-67, a lipophilic dye to visualize the M-EVs via microscopy (Fig. 2A). The vesicular nature of the EV was confirmed via transmission electron microscopy (Fig. 2B) (Supplementary fig. 1B). Nanoparticle tracking analysis assay revealed that the EVs were 100-200nm in diameter, consistent with the expected size for EVs (Fig. 2C). We used Western blotting to assess EV biomarkers and demonstrated that the EV’s were positive for both CD63 and CD81 (Fig. 2D) (Supplementary fig. 1C). These data showed the successful isolation of M-EVs.

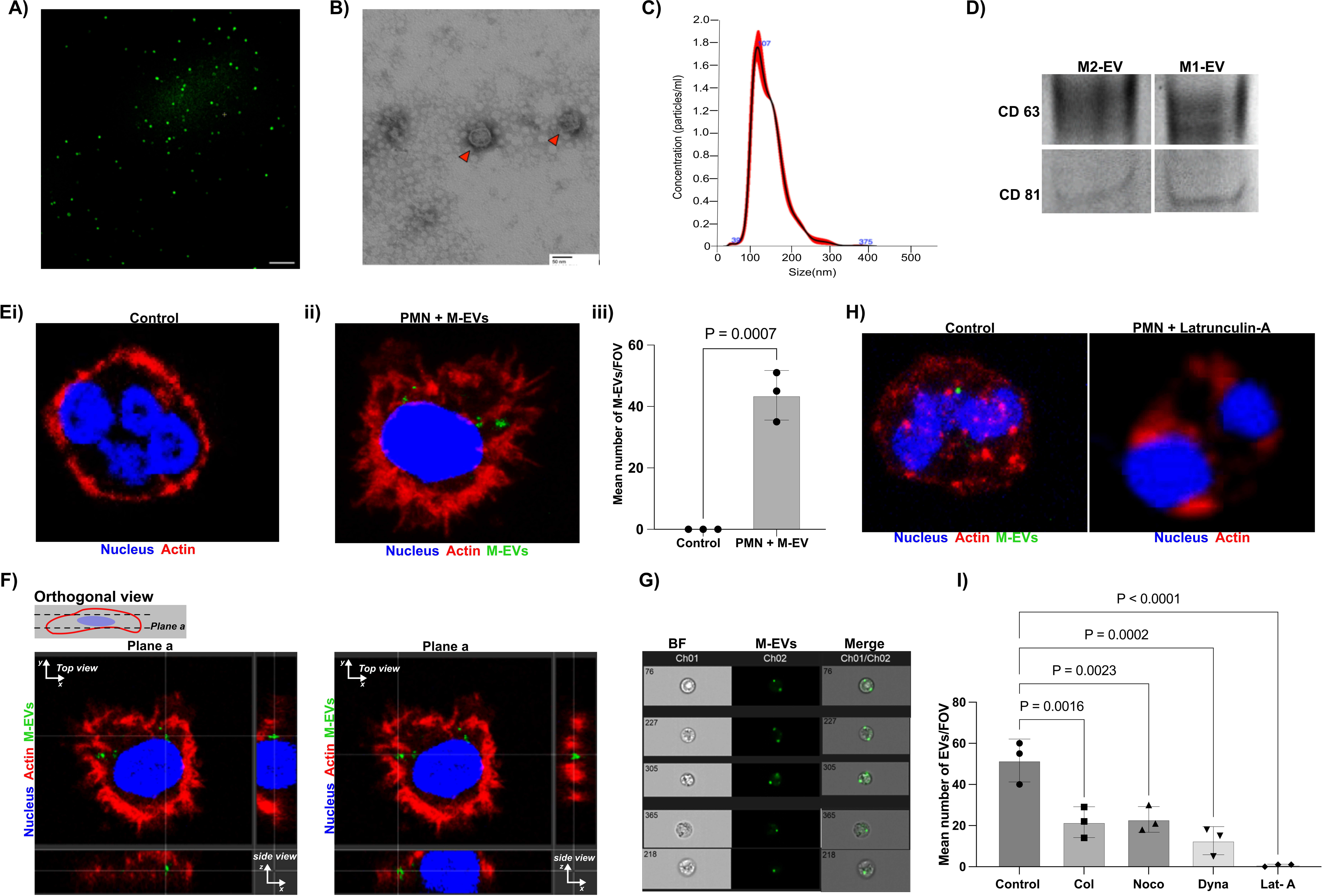

### Human serum opsonized M-EVs are internalized by neutrophils via endocytosis

There are increasing evidence that demonstrate that macrophages are involved in the regulation of tissue damage by releasing EVs with pro- or anti-inflammatory cargo to modulate inflammation in their microenvironment (*21–23*). However, it’s unclear if neutrophils interact with M-EVs at a site of inflammation, that could in turn modulate neutrophil behavior and activities during inflammation. Thus, to test whether M-EVs are internalized by neutrophils and possibly activate neutrophils during this interaction, we opsonized M-EV in autologous human serum for 30 mins and then labelled them with a membrane lipophilic PKH-67 dye. Then we co-incubated the labelled M-EV with freshly isolated neutrophils for 3-4 hours. We observed uptake of PKH-67 labelled M-EVs as confirmed by confocal microscopy (Fig. 2E & F) (Supplementary video 3) (Supplementary fig. 1D) and image stream cytometry (Fig. 2G & Supplementary. Fig. 2A-C).

Most experimental evidence suggests that EVs from various cell types are internalized into the cell via endocytosis (*24*). To test whether uptake of M-EVs into neutrophils is via an endocytic pathway, we co-incubated neutrophils with either M-EVs alone or together with a series of known endocytosis inhibitors that have previously been shown to block uptake in neutrophils(*24*). Specifically, we tested the actin filament inhibitors Latrunculin A, as actin filament inhibitors are known to block endocytosis(*25*) and the microtubule inhibitors Nocodazole and Colchicine as well the Dynamin-2 inhibitor Dynasore(*26*). Dynamin-2 is required for Caveolin-dependent endocytosis and its inhibition through Dynasore can block EV uptake(*27*). The inhibitors significantly reduced EV uptake (Fig. 2H & I), suggesting that EV internalization by neutrophils proceeds via endocytosis.

### M1-EVs and M2-EVs modulate migratory response in neutrophil during SI and nSI

M1 macrophage activity can inhibit cell proliferation and cause tissue damage while M2 macrophage activity is thought to promote cell proliferation and tissue repair (*28*). Having demonstrated that neutrophils internalize M-EVs in an active manner *in vitro*, next, we hypothesized that M1 and M2 macrophage subset derived EVs will modulate neutrophil migratory behavior differently depending on the inflammatory microenvironment.

Therefore, we introduced primary neutrophils pretreated with M1-EVs or M2-EVs into the endothelial lumen and examined their behavior using time-lapse imaging. While M-EV treatment increased PMN migration overall compared to the untreated PMN control, we observed that M1-EVs activate a larger migratory response in neutrophils compared to M2-EV during nSI. M1-EV treated neutrophil recruitment starts within the first 5-10 minutes, when they migrate across the endothelium, through a collagen matrix, to the nSI site and lasted for 1-2 hours. We observed significantly more migration of M1-EV treated neutrophils compared to the M2-EV treated group (Fig. 3A & B) (Supplementary fig. 1E) (Supplementary video 4) and (Supplementary video 5). On the other hand, using the SI model, we observed that M2-EV treated neutrophils demonstrated a greater migratory response toward the SI spheroid site compared to M1-EV treated neutrophils (Fig. 3C & D) (Supplementary fig. 1F) (Supplementary video 6) and (Supplementary video 7). Similar to the M1-EV treated neutrophils, recruitment starts within the first 10 minutes by directionally migrating through the collagen matrix to the SI site. Migration lasted for about 2 hours. Our results suggest that different macrophage subset derived EVs modulate neutrophil migratory responses differently in SI and nSI inflammatory microenvironments.

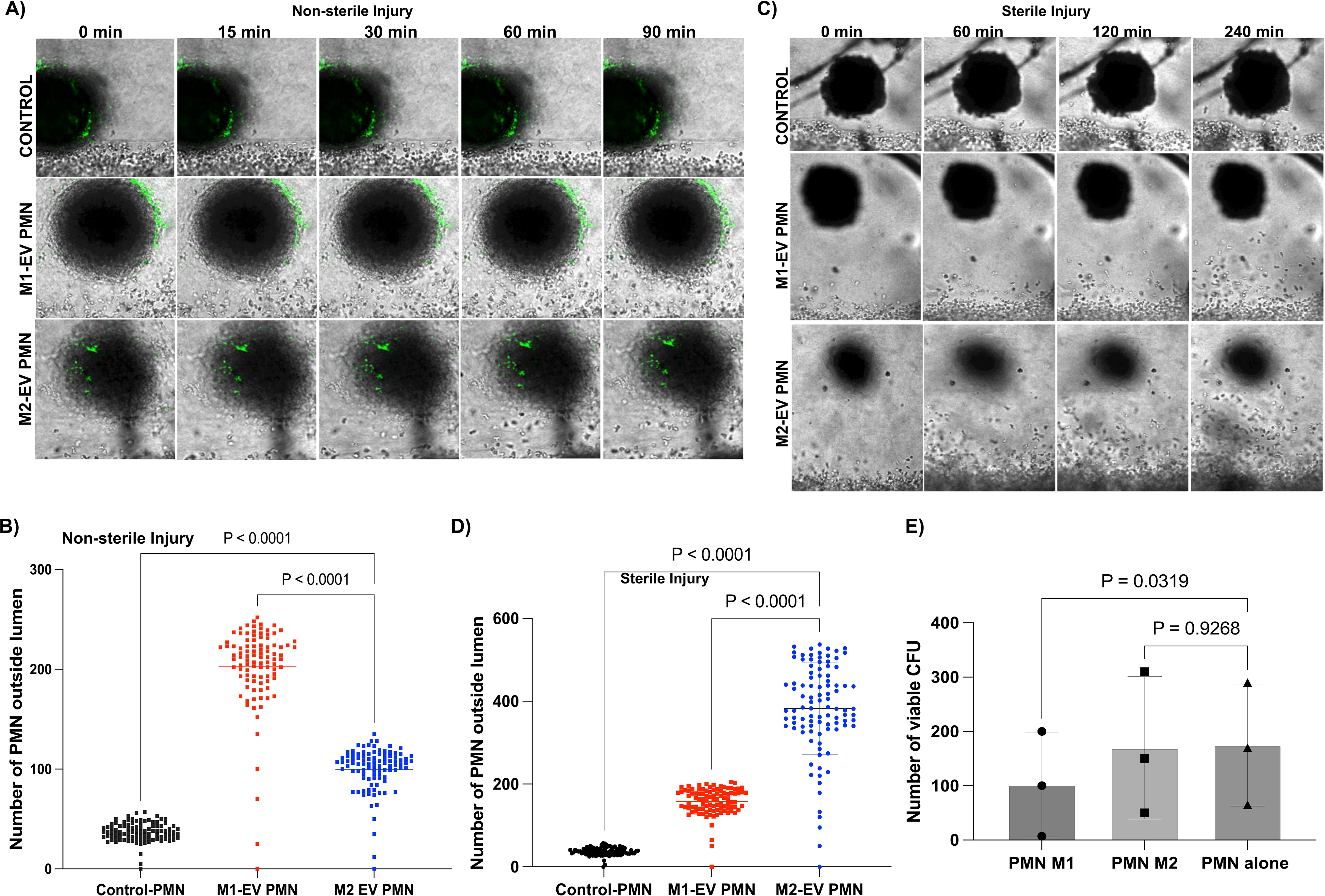

To understand the possible activities of M-EV treated PMNs at the nSI site, we carried out a colony forming unit assay (CFU) to quantify number of viable bacteria (*Staphylococcus aureus*) colonies after being incubated with M-EV treated PMNs. Briefly, neutrophils were treated with either no EVs, M1-EVs or M2-EVs for 3-4 hours. Next, the treated neutrophils were challenged with *Staphylococcus aureus* at a MOI of 1:10. After 3 hours of incubation, extracellular non phagocytosed bacteria were killed using a poorly diffusible antibiotic, gentamicin. Neutrophils were subsequently lysed in cold water and plated to count viable bacteria colonies after 24 h. The CFU assay revealed a significant increase in viable bacteria colonies in untreated neutrophils compared to M1-EV treated neutrophils, suggesting the increased capacity of M1-EV treated neutrophils to kill bacteria at the nSI site (Fig. 3E) (Supplementary fig. 3A). M2-EV treated neutrophils showed no significant difference in the number of viable bacteria compared to control.

### M2-EVs treated neutrophils migrate away from the SI site and express ICAM-1

M2 macrophages are capable of anti-inflammatory responses and repair damaged tissues(*29*). Thus, we hypothesize that M2-EVs will modulate neutrophil migratory behavior to reduce neutrophil influx to the injury site while M1-EVs will not show this effect. Using a Leica super STED resolution microscope, we monitored and tracked M1-EV treated neutrophil migration from the endothelial lumen toward the SI spheroid site. Our detailed imaging revealed that M1-EVs treated neutrophils (second panel) migrate directionally toward the SI site and randomly migrate/stay within the SI site (Fig. 4A & Bii) (Video 1).

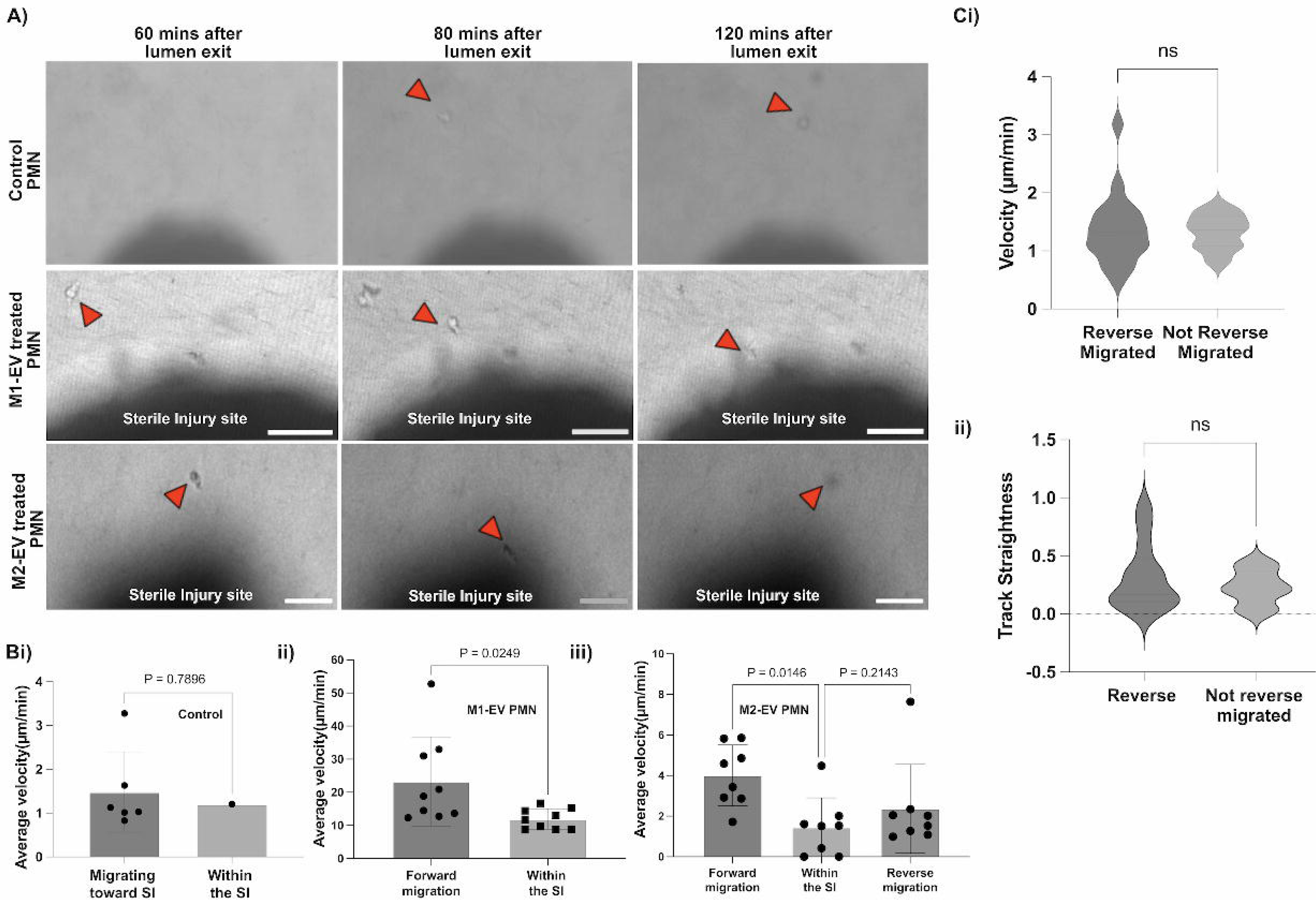

On the other hand, imaging around the SI site revealed that M2-EVs treated neutrophils (third panel) also migrated directionally toward the SI site, however, on reaching the site, they interact with the dead cells debris and immediately migrate away from the SI site (Fig. 4A) (Video 2). To identify reverse migrated neutrophils, we monitored and tracked individual cells from the point of lumen exit to and from the SI site. A neutrophil qualifies as reverse-migrated if it has travelled away from the SI site for a minimum distance (*30*). The behavior of cells during reverse migration were further described by measuring other parameters such as cell speed, cell displacement and track straightness in our model(*4*) (*30*).

Reverse migrated neutrophils (M2-EV treated neutrophils) upon reaching the SI site, either migrate toward the lumen direction or migrate randomly away into the collagen matrix. Tracking revealed that these cells show increased directional migration toward the border of the injury site, though the interaction within the injury site seems brief and was mainly around the edges of the injury sites (Fig. 4A). Control neutrophils showed no significant differences in their velocity between migration toward or within the SI site, however M1-EV treated neutrophils demonstrate significantly increased migration towards the SI compared to within the SI site. (Fig 4Bi-ii). The velocity of M2-EV treated neutrophils also increased as they migrated toward the SI site and they exhibited significantly slower velocity within the SI site (potentially due to interacting with the dead cells) (Fig 4Biii). However, on migrating away from the site, the cells displayed a slight but not significant velocity increase (Fig. 4Biii).

Next, we compared the behavior of the reverse migrated cells with the non-reverse migrated cells among the M2-EV neutrophils using the reported metrics (track velocity and straightness). We found no significant differences in the rM neutrophils compared to the non rM neutrophils (Fig. 4Ci-ii). The above results suggest that the type 2 macrophage subset could be reducing excessive influx of neutrophils to the injury site thereby contributing to the resolution of inflammation.

To confirm the phenotype of the reverse migrated cells in our MPS, we stained for the expression of markers of a reported reverse migrated neutrophil phenotype (ICAM-1^high^/CXCR-1^low^) (*5*, *31*) in our model. We observed that a sub-population of M2-EV treated neutrophils express more ICAM-1 during SI compared to other groups (Fig. 5A-C) (Supplementary Videos 8-10). Thus, further demonstrating the ability of our MPS to study the role of M2-EV in stimulating rM in primary neutrophils.

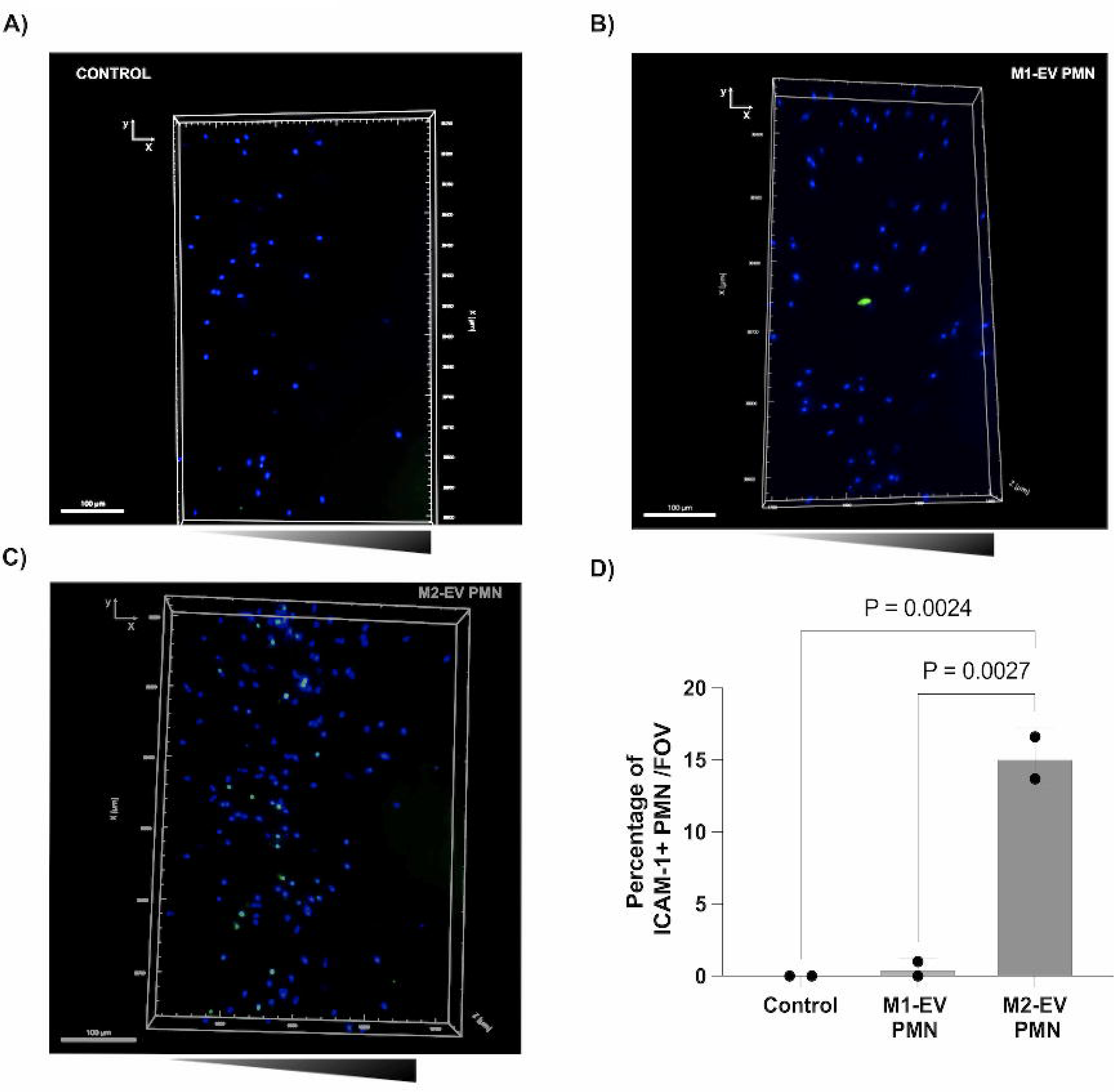

### M2-EV derived IL-8 maybe crucial to neutrophil rM *ex vivo*

Having shown that M2-EV induce neutrophils to migrate away from the SI site, we speculated that the observed behavior was most likely associated with the content of M2-EVs. Therefore, we isolated EVs from both M1 and M2 macrophages via ultracentrifugation and used a chemokine protein array blot to detect and quantify chemokine cargoes of M-EVs. Our analysis showed that IL-8 is one of the most abundant chemokines in M2-EVs compared to M1-EVs (Fig. 6A-D). We speculated that M2-EV derived IL-8 may be an important chemokine in the observed rM in primary neutrophils. To, ascertain the possible role of IL-8 in neutrophil rM, we knocked down IL-8 in human monocyte cell line (THP-1) using CRISPR-cas9. Validation of the gene knockdown was confirmed via western blot (Fig. 6E), chemokine protein array blot (Supplementary fig. 3B-C) and a migration assay (Supplementary fig. 3D).

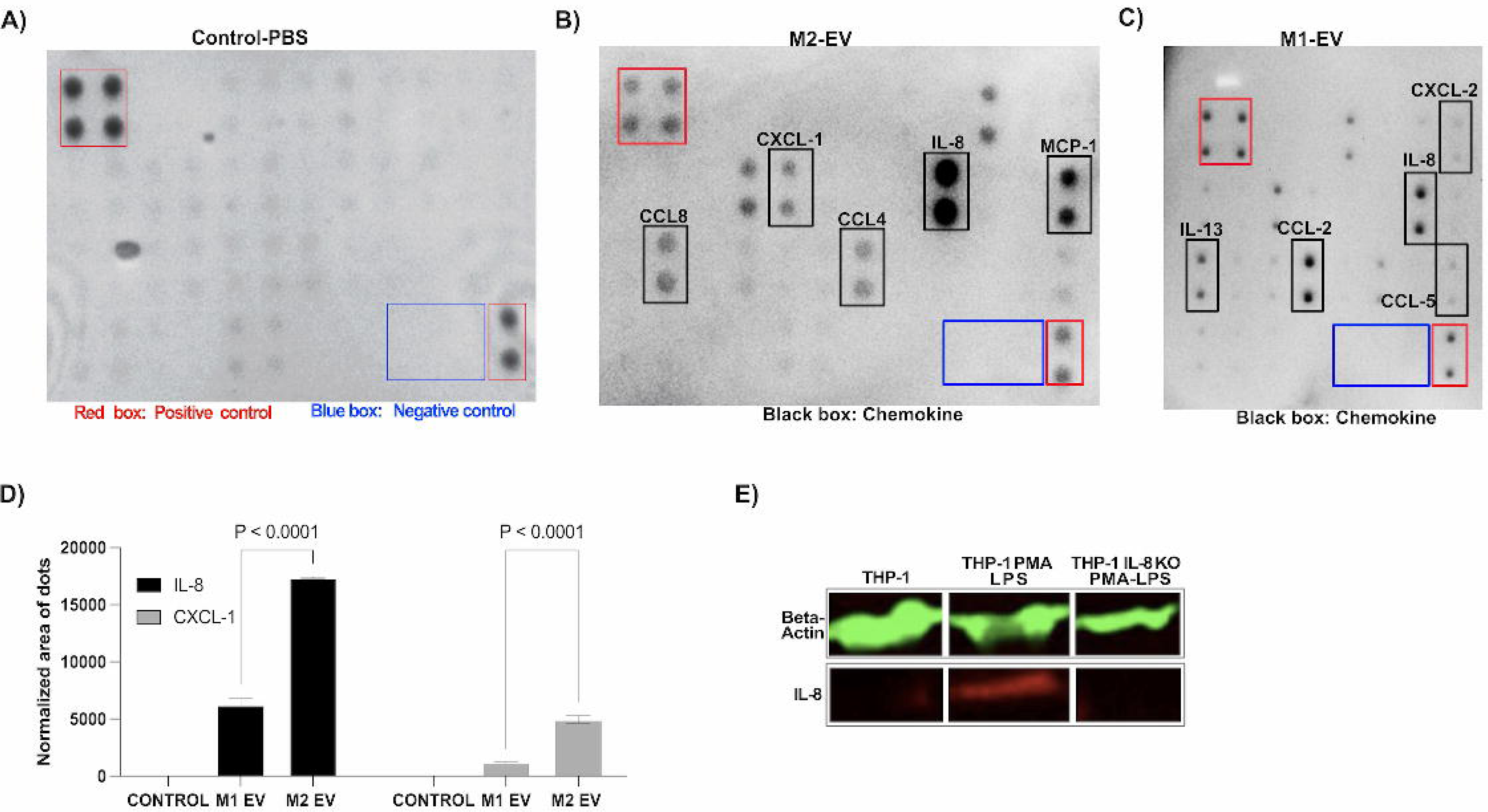

Tracking analysis revealed that neutrophils treated with IL-8 KO EVs were able to migrate through the endothelial lumen but lack directional migration toward the injury site as most cells show a random migratory pattern during SI in our model compared to M1 or M2-EV treated cells (Fig. 7Aiii) (Supplementary Fig. 3Eiii). Furthermore, we compared the behavior of the cells in terms of track straightness in each group. Importantly, our data showed that track straightness in M2-EV treated neutrophils were similar during forward migration and reverse migration (Fig. 7Bi) (Supplementary Fig. 3Eii). Similar to M2-EV treated neutrophils, track straightness was also similar in M1-EV treated neutrophils during forward migration and within the SI (Fig. 7Bii). Finally, our data showed that the IL-8 KO EV treated neutrophils showed increased track straightness compared to other groups, however with no directionality towards the SI site (Fig. 7Biii) and demonstrated no evidence of rM in our system (Fig. 7Aiii) (Supplementary Fig. 3Eiii) (Video 3).

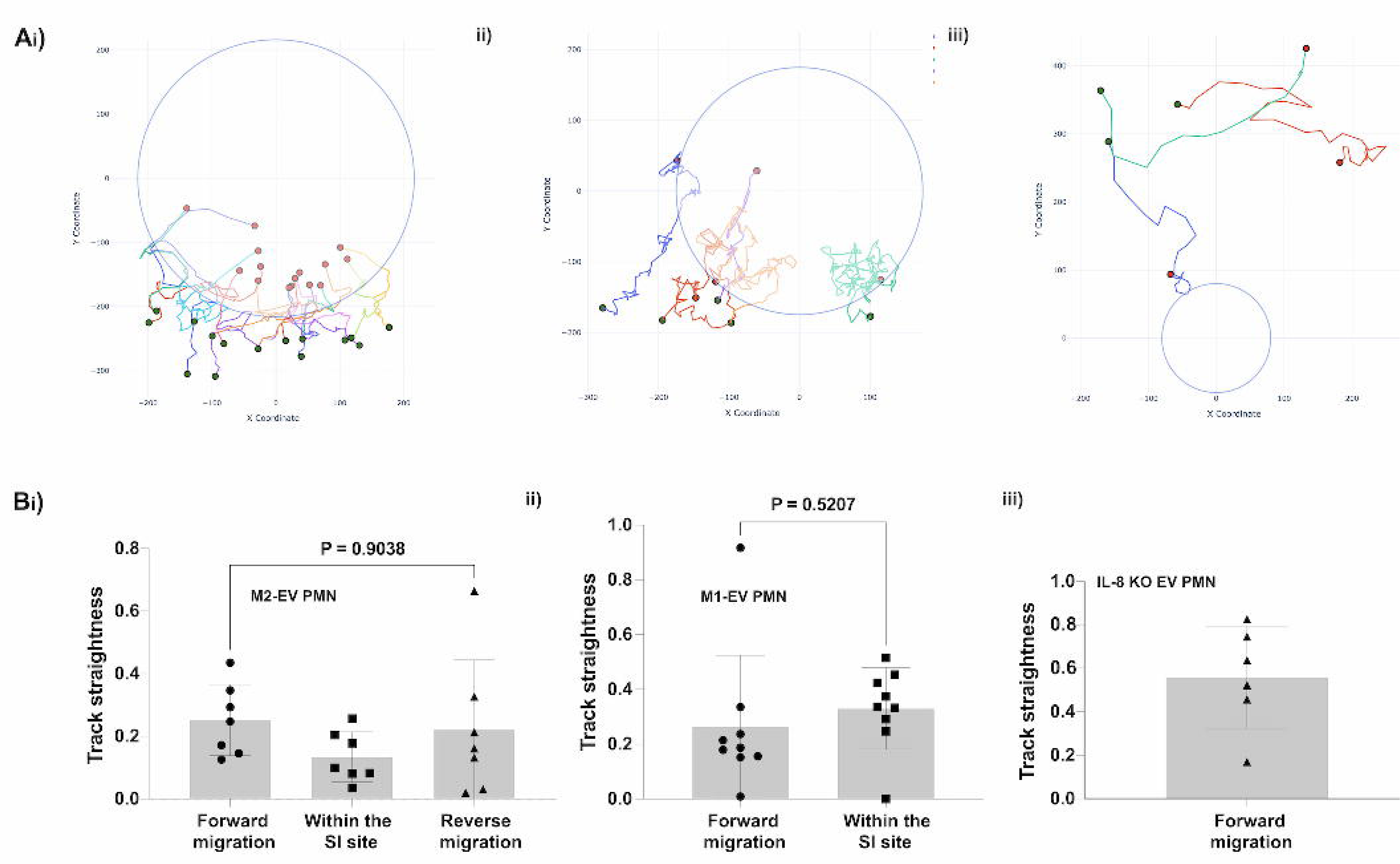

## DISCUSSION

Neutrophil reverse migration (rM) has been described as a process by which neutrophils migrate away from the inflammatory site back into the vasculature following initial infiltration, which is involved in the resolution of local inflammatory responses. rM in neutrophils has been demonstrated using animal models(*4*, *17*). However, evidence of rM in human primary neutrophils are still lacking. The present study aimed to demonstrate rM in human primary neutrophils and explore the possible mechanisms involved in human primary neutrophil rM. By using an MPS incorporated with human cells, we demonstrated that the interaction between M-EVs and neutrophils that could lead to the resolution of inflammation via neutrophil rM.

In this study, we demonstrate a new mechanism underlying rM, whereby M2-EV-derived IL-8 elicits a distinct migratory pattern (rM) in neutrophils using our SI MPS. We identified a reverse migrated neutrophil as one that has travelled away from the SI site for a minimum distance. We then quantified the migratory behavior of the reverse migrated neutrophils using metrics: (a) track speed and (b) track straightness. By enabling quantitative studies of the interaction between neutrophils and M-EVs in our platform, our MPS platform could create opportunities to design EV related therapeutic agents that target diseases related to excessive influx of neutrophils to the injury site.

The key features of our MPS enable characterization of neutrophils migration both toward and away from the SI site. Neutrophils extravasate out of the lumen into the ECM following inflammatory signal gradients generated by a SI spheroid adjacent to the lumen. The self-generating inflammatory signal source (SI spheroid) is endogenous and mimics injury site architecture. While its known that neutrophil migratory behavior may be dependent on the context of the local microvascular and tissue environment (*32*), this important pathophysiological issue is not addressed in previous studies using traditional assays such as Dunn and Zigmond chambers(*33*), where cells migrate on flat surfaces and other microfluidic platforms(*11*), where cells migrate in straight channels. In our MPS, neutrophils migrate in a more physiological microenvironment, thus recapitulating environmental cues that neutrophils encounter *in vivo* as they transmigrate from the endothelium and within the extracellular matrix, where neutrophils move between cells within tissues(*4*, *34*). In addition, previous work from our group has shown that neutrophils live longer, ∼18-24 hours in our MPS platform, thus enabling us to study and monitor neutrophil migratory behavior for longer period (*35*)

The novelty of our MPS relies on incorporating an inflammatory injury site, primary immune cells and immune cell-derived vesicles within scaffolding extracellular matrices. Using the SI MPS, we studied how M-EVs modulate neutrophil behavior at the injury site by introducing M-EV pre-treated primary neutrophils directly into the endothelial lumen. This approach allowed neutrophils to transmigrate through the endothelial lumen prior to migrating to the inflammatory stimulus mimicking the *in vivo* process. Using live cell microscopy, we demonstrated rM driven by neutrophil-M-EV crosstalk in primary neutrophils *in vitro*. It is well known that EVs may function as carriers, as they are made up of a lipid bilayer and contain RNA and proteins(*19*). Previous studies have demonstrated that EVs can mediate cell–cell communication by transferring RNAs or proteins into target cells(*36*).

Neutrophils are rapidly recruited to infection sites, where they mediate effective clearance. It is not clear whether M-EVs modulate neutrophils behavior at the site of inflammation. One of the key findings enabled by our MPS is that it demonstrates the specific interaction between neutrophils and M-EVs via phagocytosis that in turn modulate neutrophil migratory patterns during SI and nSI at a high spatiotemporal resolution. This interaction would be challenging to capture *in vivo* because of the complexities involved in *in vivo* imaging.

Chemokines play a key role in the recruitment of neutrophils. Macrophages are an important source of chemokines during inflammation. Therefore, we chose macrophages as the initiating and modulating factor of neutrophil chemotaxis as tissue macrophages have been demonstrated to play an important role in the synthesis of CXCL2 and CXCL1 during neutrophil recruitment (*37*). IL8 is an important chemokine that mediates neutrophil migration and function in inflammation(*38*, *39*). Importantly, using our MPS, we showed that IL-8 (M2-EV derived IL-8), elicits neutrophil migration both towards and away from the SI site. While the chemo-attractive and chemo-repulsive actions of IL-8 have been previously reported(*11*, *40*) at high IL-8 concentrations, we find that the forward and reverse migration patterns are not preserved when IL-8 levels were reduced (IL-8 KO M2-EV).

The differences between our findings and previous findings may be due to differences in the cell microenvironment due to experimental settings, such as ECM versus flat glass surfaces, transmigration through the endothelial lumen, and physiological signals from vs chemokines. This possibility is supported by the dissimilarities in the neutrophil migration speed, 3-5 μm min^−1^ in our MPS versus 18.6–21.5 μm min^−1^ in straight channels(*11*). This IL-8 driven forward and reverse migration pattern would be difficult to capture in a Boyden chamber or in a transwell assay, where cells initially migrating through the membrane are not able to reverse direction.

Furthermore, our MPS allowed us to study the behavior of the reverse migrated cells by studying migratory parameters such as track velocity and straightness before and during reverse migration. We show that track straightness during forward and reverse migration in M2-EV treated cells is similar thus, suggesting similar mechanism in both migration modes (*30*). Though IL8-KO EV treated neutrophils showed more track straightness compared to other groups, most of the cells lack directionality toward the SI site.

One of the study limitations is that neutrophil rM is a relatively rare process with only a subset of the cells possessing the ability to undergo this process. Therefore, this restricts the number of cells available for study in the MPS. Other limitations include that we did not examine the effect of other chemokine cargoes (CCL2 and CXCL-1) of M2-EVs on neutrophil migratory response. Further, our model lacks the ability to track the destination of the reverse migrated neutrophils (for example, the lungs, liver and bone marrow). Future experiments could mimic these organs in our SI MPS. Thirdly, RNA sequencing analysis and examination of additional immunomodulatory factors could help provide a more in-depth molecular analysis of the biological processes activated in neutrophils by M2-EVs driving rM during SI in our model.

The primary objectives of this work were to create an MPS that can investigate specific interactions between immune cells (such as neutrophil macrophage interaction via EVs) at the site of injury (such as rM) that can be challenging to study using animal models. Our approach to generating 3D geometries of blood vessels and focal inflammatory injury sites within biomimetic ECM scaffolds could answer questions that have so far been difficult to address such as isolating reverse migrated human primary neutrophils and understanding the genetic pathway in rM neutrophils via single cell RNA sequencing. Combined with the ability to dynamically visualize and explore immune responses to SI inflammation at a high spatiotemporal resolution, our model may have potential for discovery of immune cell derived EV therapies for non-healing wounds.

## ACKNOWLEDGMENTS

This work was supported by the University of Wisconsin Carbone Cancer Center, Cancer Center Support Grant NIH P30CA014520. This work was also supported by the National Institutes of Health NIH R01AI134749 to D.B., and the National Institutes of Health NIH U24AI152177 to D.B. and S.K., and the Jubilaumsstiftung von Swiss Life (project number: 1350) to K.A.B. We thank Alice Golubiewski in the MMB Lab for arranging whole blood collection from healthy donors. We thank the University of Wisconsin Carbone Cancer Center Flow Cytometry Laboratory for the use of its facilities and services. We also want to thank Dr J.D Sauer for the gift of the *S.A gfp* strain. We would like to thank Dr David Bennin for training and use of the Lonza Nucleofector. Finally, we thank the University of Wisconsin Optical Imaging Core for their support with the confocal imaging and use of the imaging facilities.

## AUTHOR CONTRIBUTIONS

K.A.B designed and performed experiments and did data analysis with assistance from A.A., B.F.O., D.J.B., and S.C.K. B.F.O and W.S.P performed all the western blot technique and analysis. A.A carried out the python coding and analysis and generation of figure 1D. K.A.B., B.F.O., A.A., W.S.P., D.J.B., and S.C.K. wrote and reviewed the manuscript.

