## Supplemental Video Legends for "A microphysiological system of sterile injury demonstrates neutrophil reverse migration via macrophage-derived extracellular vesicle crosstalk" (from another version of this preprint: version 1, 2024-12-30)

**SUPPLEMENTARY VIDEOS LEGEND**

**Sterile injury micro-physiological system demonstrates neutrophil reverse migration via macrophage derived extracellular vesicles interaction *in vitro***

**Kehinde Adebayo Babatunde^1^, Babatunde Fatimat Oluwadamilola^2^, Adeel Ahmed^1^, Wilmara Salgado-Pabon^2^, David J Beebe^1,3,4^ and Sheena C Kerr^4*^.**

**^1^Department of Pathology & Laboratory Medicine, University of Wisconsin, Madison, WI, USA.**

**^2^Department of Veterinary Medicine, University of Wisconsin, Madison, WI, USA.**

**^3^Department of Biomedical Engineering, University of Wisconsin-Madison, Madison, WI 53715, USA.**

**^4^Carbone Cancer Center, University of Wisconsin, Madison, WI, USA.**

**Video 1. Primary neutrophils treated with M1-EVs migration toward SI site.**

Primary neutrophils pre-treated with M1-EVs show migratory response toward the SI site. They either migrate within the SI site or stay within the site on getting to the injury site. Neutrophils are in brightfield. The time interval between frames is 3 minutes. Scale bar is 100 μm.

**Video 2. Primary neutrophils treated with M2-EVs migration toward SI site.**

Primary neutrophils pre-treated with M2-EVs show migratory response toward the SI site. They migrate toward the S1 site and reverse migrate away on getting to the injury site. Neutrophils are in brightfield. The time interval between frames is 3 minutes. Scale bar is 100 μm.

**Video 3. Primary neutrophils treated with IL-8 KO EVs migration toward SI site.**

Primary neutrophils pre-treated with IL-8 KO EVs show migratory response but lack directionality toward the SI site. Neutrophils are in brightfield. The time interval between frames is 3 minutes. Scale bar is 100 μm.

**Supplementary Video S1. 3D conformation of the HUVEC lumen.** 3D rendering of the HUVEC seeded lumen. The nucleus of the HUVEC is stained with DAPI (blue) and calcein (red). Scale bar is 150 μm.

**Supplementary Video S2. 3D conformation of the spheroidal nonsterile injury site.** 3D rendering of the nSI site. The nucleus of the dermal fibroblast is stained with Hoechst (blue), Actin (red) and *S. aureus* (GFP). Scale bar is 50 μm.

**Supplementary Video S3. 3D conformation of the M-EV uptake by primary neutrophil.**

Primary neutrophils were incubated with M-EVs and immunofluorescent staining was carried out to confirm uptake. The nucleus of the primary neutrophil is stained with Hoechst (blue), Actin (red) and M-EV (PKH-67). Scale bar is 20 μm.

**Supplementary Video S4. Primary neutrophils treated with M1-EVs migration toward nSI site.** Primary neutrophils pre-treated with M1-EVs show migratory response toward the nSI site. Neutrophils are in brightfield and *S. aureus* (GFP). The time interval between frames is 3 minutes. Scale bar is 50 μm.

**Supplementary Video S5. Primary neutrophils treated with M2-EVs migration toward nSI site.** Primary neutrophils pre-treated with M2-EVs show migratory response toward the nSI site. Neutrophils are in brightfield and *S. aureus* (GFP). The time interval between frames is 3 minutes. Scale bar is 50 μm.

**Supplementary Video S6. Primary neutrophils treated with M1-EVs migration toward SI site.** Primary neutrophils pre-treated with M1-EVs show migratory response toward the SI site. Neutrophils are in brightfield. The time interval between frames is 3 minutes. Scale bar is 50 μm.

**Supplementary Video S7. Primary neutrophils treated with M2-EVs migration toward SI site.** Primary neutrophils pre-treated with M2-EVs show migratory response toward the SI site. Neutrophils are in brightfield. The time interval between frames is 3 minutes. Scale bar is 50 μm.

**Supplementary Video S8. Expression of ICAM-1 on primary neutrophils alone during SI.** Expression of ICAM-1 in primary neutrophils alone during migratory response toward the SI site. The nucleus of the neutrophils was stained with Hoechst and ICAM-1 (Alexa flour 488). Scale bar is 100 μm.

**Supplementary Video S9. Expression of ICAM-1 on primary neutrophils treated with M1-EVs during SI.** Expression of ICAM-1 in primary neutrophils treated with M1-EVs during migratory response toward the SI site. The nucleus of the neutrophils was stained with Hoechst and ICAM-1 (Alexa flour 488). Scale bar is 100 μm.

**Supplementary Video S10. Expression of ICAM-1 on primary neutrophils treated with M2-EVs during SI.** Expression of ICAM-1 in primary neutrophils treated with M2-EVs during migratory response toward the SI site. The nucleus of the neutrophils was stained with Hoechst and ICAM-1 (Alexa flour 488). Scale bar is 100 μm.
