## Supplementary Figure Legends for "A microphysiological system of sterile injury demonstrates neutrophil reverse migration via macrophage-derived extracellular vesicle crosstalk" (from another version of this preprint: version 1, 2024-12-30)

**Sterile injury micro-physiological system demonstrates neutrophil reverse migration via macrophage derived extracellular vesicles interaction *in vitro***

**Kehinde Adebayo Babatunde^1^, Babatunde Fatimat Oluwadamilola^2^, Adeel Ahmed^1^, Wilmara Salgado-Pabon^2^, David J Beebe^1,3,4^ and Sheena C Kerr^4*^.**

**^1^Department of Pathology & Laboratory Medicine, University of Wisconsin, Madison, WI, USA.**

**^2^Department of Veterinary Medicine, University of Wisconsin, Madison, WI, USA.**

**^3^Department of Biomedical Engineering, University of Wisconsin-Madison, Madison, WI 53715, USA.**

**^4^Carbone Cancer Center, University of Wisconsin, Madison, WI, USA.**

**FIGURE 1:** A) Representative images of monocyte differentiated macrophage polarization to M1 macrophage (showing the expression of CD 80 marker) and M2 macrophage (showing the expression of CD 163 marker) *in vitro*. Scale bar is 20µm. B) Electron microscopy image of M-EVs. Mag: 66000x. Scale bar: 50nm. Red arrow points to the M-EV. C) WB of M-EV markers showing the protein ladder. D) Immunofluorescence images showing neutrophil with M-EVs; Neutrophil nuclei (DAPI-Blue), Actin filament (Phalloidin-Red) and M-EVs (PKH-67-Green). Scale bar: 10µm. Migratory dynamics and number of neutrophils outside the lumen during E) nSI and F) SI.

**FIGURE 2:** A) Image stream cytometry gating analysis showing population of neutrophil with M-EVs and without M-EVs. B) Image stream software analysis of neutrophils showing different counts of M-EVs being uptaken by neutrophils.

**FIGURE 3:** A) Representative image of viable bacteria agar plate after incubation with control, M1-EV PMN and M2-EV PMN. Chemokine array analysis image showing IL-8 cargo in B) Control THP-1 cells and C) IL-8 THP-1 KO cells. D) Graph showing number of neutrophils that responded to THP-1 like macrophage cells and IL-8 KO THP-1 like macrophage cells (chemotaxis assay). E) Representative heatmap of the migratory tracks in i) M1-EVs treated neutrophils, ii) M2-EV treated neutrophils and iii) IL8 KO EV treated neutrophils toward the SI site. Black circles represent the boundary of the SI site.
